# Dihydroceramide desaturase regulates the compartmentalization of Rac1 for neuronal oxidative stress

**DOI:** 10.1101/2020.06.01.128579

**Authors:** Fei-Yang Tzou, Tsu-Yi Su, Yu-Lian Yu, Yu-Han Yeh, Chung-Chih Liu, Shu-Yi Huang, Chih-Chiang Chan

## Abstract

Disruption of sphingolipid homeostasis has been shown to cause neurological disorders. How specific sphingolipid species modulate the pathogenesis remains unknown. The last step of sphingolipid *de novo* synthesis is the conversion of dihydroceramide to ceramide catalyzed by dihydroceramide desaturase (human DEGS1; *Drosophila* Ifc). Loss of *ifc* leads to dihydroceramide accumulation and oxidative stress, resulting in photoreceptors degeneration, while *DEGS1* variants were associated with leukodystrophy and neuropathy. Here, we demonstrated that *ifc* regulates Rac1 compartmentalization in fly photoreceptors and further showed that dihydroceramide alters the association of active Rac1 to membranes mimicking specific organelles. We also revealed that the major source of ROS originated from Rac1 and NADPH oxidase (NOX) in the cytoplasm, as the NOX inhibitor apocynin ameliorated the oxidative stress and functional defects in both fly *ifc*-KO photoreceptors and human neuronal cells with disease-associated variant *DEGS1*^*H132R*^. Therefore, *DEGS1*/*ifc* deficiency causes dihydroceramide accumulation, resulting in Rac1 translocation and NOX-dependent neurodegeneration.

**Graphical Abstract:** **A** *DEGS1/ifc* converts dihydroceramide to ceramide in neuronal cells, and the endolysosomal NOX complex is not activated.
**B** Dihydroceramide accumulates without functional *DEGS1/ifc* and causes alterations in membrane microdomains and recruits active Rac1 to endolysosomes. The activation of endolysosomal Rac1-NOX complex elevates cytosolic ROS levels, causing neurodegeneration.

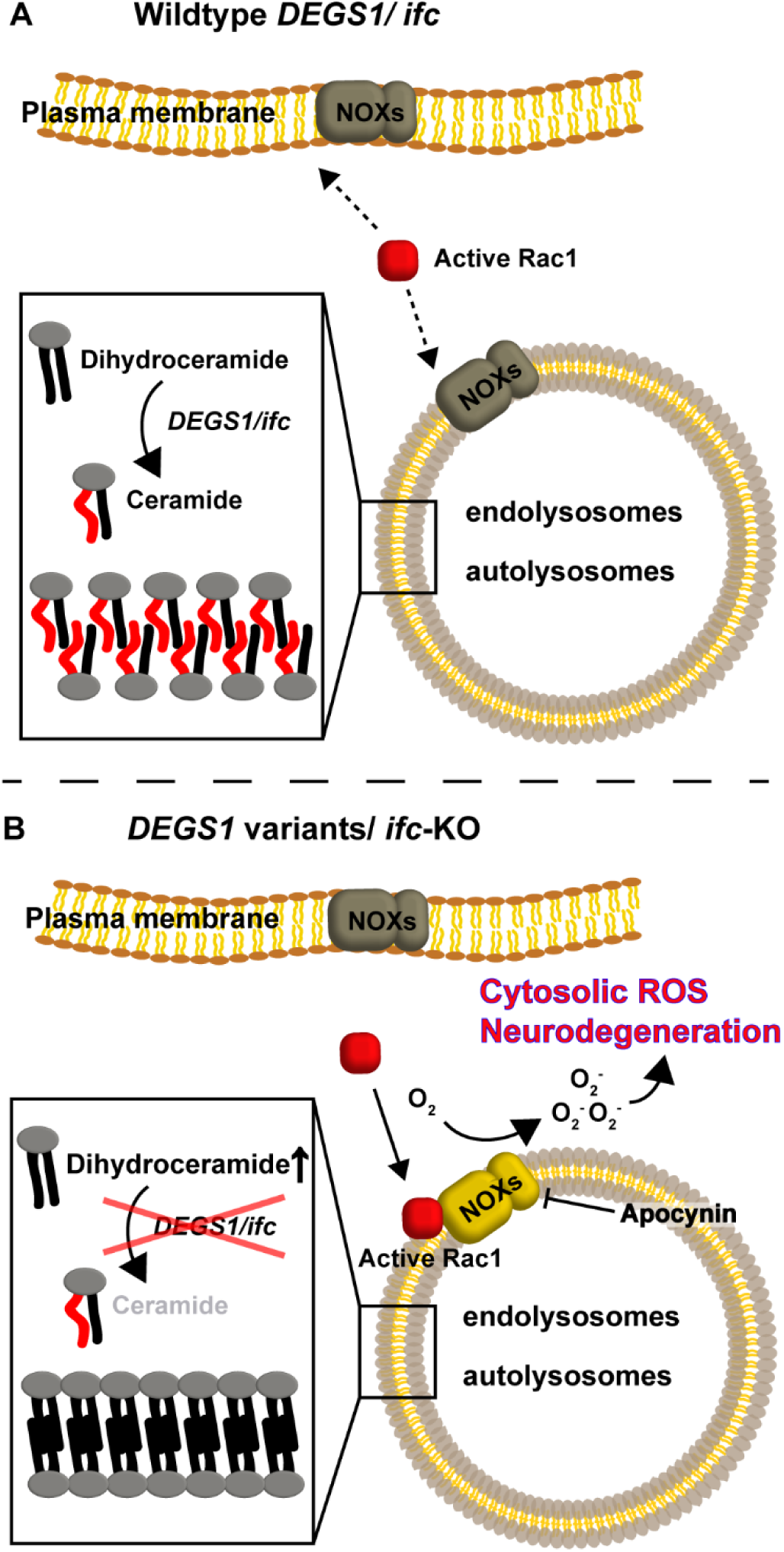

**In Brief (eTOC blurb):** Deficiency in dihydroceramide desaturase causes oxidative stress-mediated neurological disorders. Tzou and Su et al. show that increased dihydroceramide causes the relocalization of active Rac1, whilst inhibition of the Rac1-NOX ameliorates the oxidative stress and neuronal defects. NOX inhibitor apocynin may provide new direction of treatments for patients with *DEGS1* variants.

**Highlights:**

- Deficiency in dihydroceramide (dhCer) desaturase induces cytoplasmic ROS elevation
- dhCer alters the binding of active Rac1 to reconstituted organelle membranes
- Active Rac1 is enriched in endolysosomes in *ifc*-KO neurons for ROS genesis
- Rac1-NADPH oxidase elicits ROS, degenerating leukodystrophy-related neuronal cells

## Introduction

Sphingolipids are major components of neuronal membrane bilayers and are pivotal to the function of neurons. The last step of the *de novo* synthesis pathway is catalyzed by dihydroceramide desaturase (DEGS) to convert dihydroceramide to ceramide. Neuronal dihydroceramide is much less abundant than ceramide (Qin et al., 2010). For a long time, dihydroceramide was deemed inert because the exogenous short-chain analogs did not cause any noticeable defects (Hunot et al., 1997) and the physiological role of dihydroceramide was largely overlooked. In recent years, increased levels of dihydroceramide in cerebrospinal fluid have been observed in Alzheimer’s disease (AD) (Fonteh et al., 2015), and a high level of plasma dihydroceramide could predict the decline of cognitive function in AD patients in a longitudinal study (Mielke et al., 2011). Altered levels of dihydrosphingolipids were also noted in a mouse model of Huntington’s disease (Pardo et al., 2017). These findings inferred that the level of dihydroceramide may be important for neural function. Previously, we found that the *Drosophila* dihydroceramide desaturase, *infertile crescent* (*ifc*), is required for neuronal maintenance, as neuron-specific knockout of *ifc* caused the activity-dependent degeneration of the photoreceptors (Jung et al., 2017). Notably, the defects of *ifc*-KO flies were rescued by overexpression of a human homolog, *DEGS1*, suggesting functional conservation between fly and human. In 2019, three independent investigations described the association of *DEGS1* loss-of-function variants with leukodystrophy and systemic neuropathy (Dolgin et al., 2019; Karsai et al., 2019; Pant et al., 2019). Despite the clinical descriptions in patients and phenotypic characterization of the fly model, the pathophysiological role of *ifc*/*DEGS1* and dihydroceramide in the nervous system remains poorly understood.

A high level of reactive oxygen species (ROS) is a hallmark in *ifc*-KO photoreceptors and patient fibroblasts with *DEGS1* variants (Jung et al., 2017; Pant et al., 2019). An elevated level of ROS in *ifc*-KO photoreceptors causes morphological and functional defects that can be ameliorated by pan-antioxidant AD4 (Jung et al., 2017), but the source of ROS was unknown. The major sources of ROS include mitochondrial electron transport chain, NADPH oxidases (NOXs), xanthine oxidases, peroxisomal acyl-coenzyme A oxidase 1, and nitric oxide synthase, all of which have been linked to neuronal oxidative stress in different pathological contexts (Angelova and Abramov, 2018; Chung et al., 2020; Dawson and Dawson, 2018; Honorat et al., 2013; Sorce et al., 2017). Of note, pan-antioxidants therapies were promising in preclinical models but had no significant effects in clinical trials. Targeted antioxidants with subcellular precision might provide better therapeutic potential. For instance, mitochondrial-targeted antioxidants have shown protective effects in preclinical models and undergone clinical trials to treat diseases manifesting mitochondrial oxidative stress (Snow et al., 2010; clinicaltrial.gov, NCT00329056, NCT04267926, NCT03514875). Therefore, identifying wherein subcellular ROS is generated and regulated in neuronal cells may reveal the mechanistic cause of oxidative stress-induced neurodegeneration hence provides a direction for potential therapeutics.

Proteins were recruited as signaling molecules to membrane rafts enriched in cholesterol and sphingolipids, and these molecules may exhibit binding preference depending on the degree of lipid packing for protein-membrane associations (Kulakowski et al., 2018; Resh, 2004). Both *ifc*-KO flies and *DEGS1*-deficient patient fibroblasts manifested a lipid repertoire with increased dihydroceramide-to-ceramide ratio (Jung et al., 2017; Pant et al., 2019). Compared to ceramide-enriched membranes, biophysical studies show that dihydroceramide-enriched membranes are more condensed, contained less lipid packing gaps, and associate more favorably with sphingomyelin probably due to stronger intermolecular interactions (Kinoshita et al., 2020). The difference between dihydroceramide and ceramide in their intermolecular interactions with other lipids could affect the composition and packing of membrane rafts (Alanko et al., 2005). Whether dihydroceramide accumulation alters the interaction between membrane and membrane-associated proteins remained unexplored.

Here, we found that the knockout of *ifc* translocated active Rac1 to endolysosomal compartments as a result of dihydroceramide accumulation. We first identified the major source of ROS in *ifc*-KO eyes, and found the activation of Rac1-NOX responsible for the cytoplasmic ROS production in *ifc*-KO photoreceptors. Consistently, the NOX inhibitor apocynin reduced the oxidative stress in both *ifc*-KO photoreceptors and *DEGS1*^*H132R*^ SH-SY5Y cells and ameliorated the functional loss of *ifc*-KO eyes. Our colocalization study revealed an increased interaction between active Rac1 and Rab7 in *ifc*-KO photoreceptors. To reveal the molecular mechanism, our liposome binding assay demonstrated that dihydroceramide reduced the binding of active Rac1 to membrane raft but enhanced its association with autophagosomes. This study highlights that the subcellular compartmentalization of Rac1 is mediated by altered dihydroceramide levels and that the downstream NOX-dependent ROS genesis caused degeneration in *DEGS1* and *ifc* deficient neurons.

## Results

### Clearance of *ifc*-mediated ROS by cytoplasmic SOD and Catalase

We have previously reported that *ifc*-KO eyes manifested ROS accumulation and subsequent neurodegeneration (Jung et al., 2017). To probe the intracellular origin wherein ROS is generated in *ifc*-KO photoreceptors, we overexpressed the cytoplasmic and mitochondrial superoxide dismutases, *SOD1* and *SOD2*, respectively, and examined their protective effects against the activity-dependent neurodegeneration. The level of ROS significantly decreased upon overexpression of *SOD1* but not *SOD2* in *ifc*-KO photoreceptors (Figures 1A-E), indicating that ROS were present mainly in the cytoplasm. Further, overexpression of *SOD1* rescued the functional and morphological defects of *ifc*-KO eyes as indicated by rhabdomere counts (Figures 1F-J), and the ERG measurement (Figures 1K-O), respectively, suggesting that the cytosolic ROS led to the dysfunction of photoreceptors. Consistently, overexpression of *catalase*, the antioxidant gene downstream of *SOD1*, also rescued photoreceptor degeneration (Figure S1). While exogenous *SOD2* expression did not ameliorate the accumulation of ROS (Figures 1D) or irregular morphology (Figures 1I), it restored the function of *ifc*-KO photoreceptors (Figures 1N), implying a minor role of mitochondrial superoxide in the *ifc*-KO neurodegeneration. To further pinpoint the site of ROS genesis, we performed a clonal analysis with the reduction-<u>o</u>xidation sensitive GFP (roGFP) biosensor tethered to the cytoplasm and mitochondrial matrix in *ifc*-KO eyes (Albrecht et al., 2011). The oxidization of roGFP by ROS reduces its fluorescent intensity at 488 nm excitation. Prior to the degeneration of *ifc*-KO photoreceptors, we found that the mutant clones exhibited lower cyto-roGFP intensity than its wild-type sister clones labeled with RFP (Figures 1P-P’’), indicating higher levels of ROS in the cytoplasm of the mutant clones. In contrast, no observable difference in the intensity of the mito-roGFP was seen between *ifc*-KO and control photoreceptors (Figures 1Q-Q’’). Taken together, the protective effect of *SOD1* overexpression and the lowered intensity of cyto-roGFP fluorescence in mutant photoreceptors showed that the ROS accumulation in *ifc*-KO photoreceptors mainly originated in the cytoplasm.

**Figure 1.**
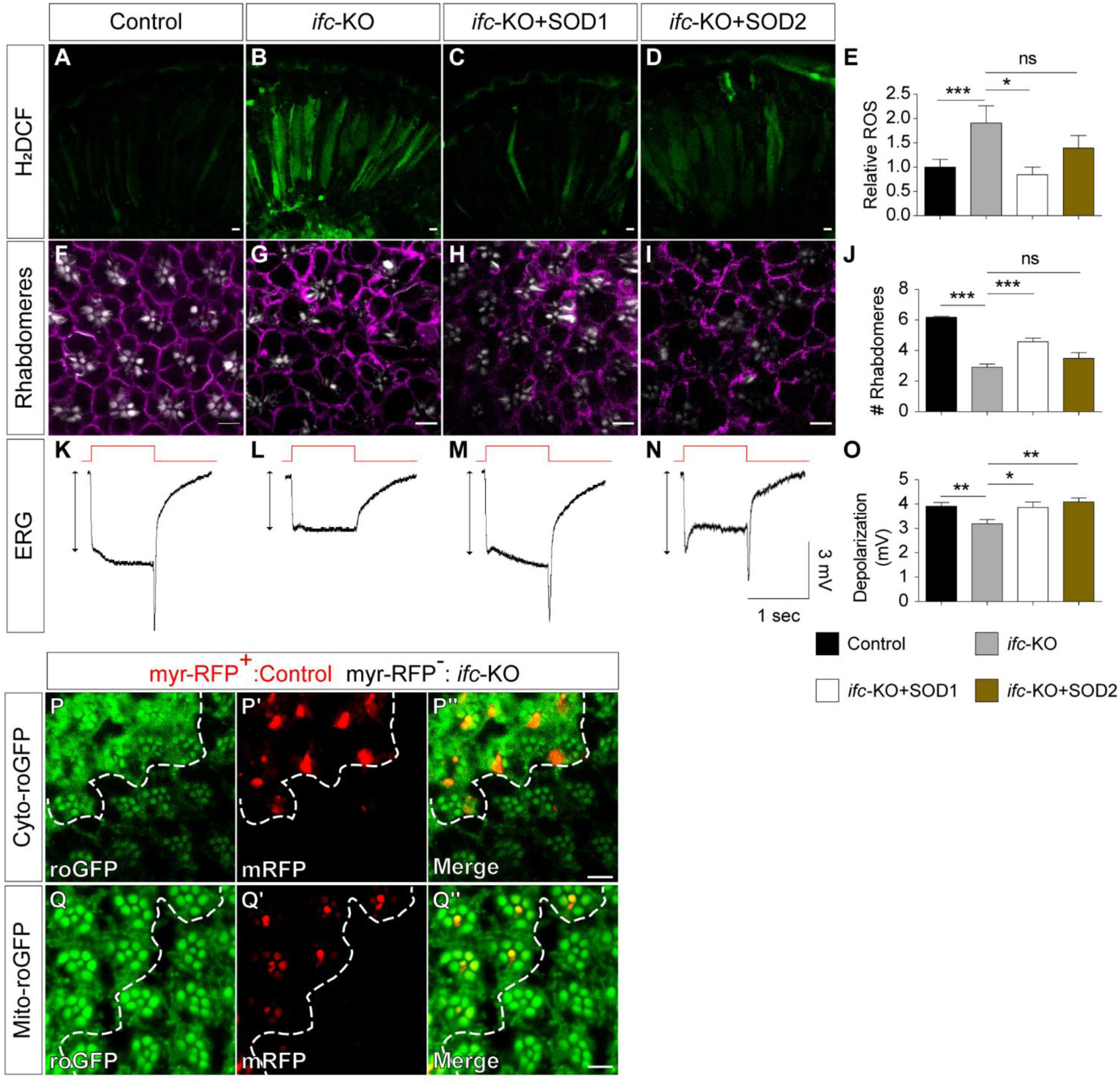
Cytoplasmic ROS accumulation in *ifc*-KO photoreceptors. (A-E) Photoreceptors longitudinal sections (A-D) stained with the ROS fluorescent probe H2DCF (green) after 3 days of light exposure. The relative ROS levels were quantified in (E) with the intensity of control photoreceptors set to 1. (F-J) Photoreceptor cross-sections (F-I) stained with Phalloidin (white; rhabdomere marker) and Na+-K+-ATPase (purple; cell membrane marker) after 5 days of light exposure. The average number of rhabdomeres per ommatidia was quantified in (J). (K-O) Representative ERG traces (K-N) of flies upon light stimulations. Double-ended arrows indicate depolarization. Flies were exposed to light for 5 days before ERG recordings. The average depolarization amplitudes (mV) of adult fly eyes was quantified in (O). (P-Q’’) *ifc*-KO mosaic eye clones (*eyflp; GMR-myr*-RFP, *FRT40A/ifc*-KO, *FRT40A*; *GMR*-Gal4/*UAS*-cyto (or mito)-roGFP) with *GMR*-driven Cyto-roGFP (P) or Mito-roGFP (Q) expression (green). Control photoreceptors were marked with *GMR*-myr-RFP (red), separated by the dashed line from *ifc*-KO photoreceptors lacking RFP signals. Scale bars: 5 μm. Error bars represent SEM; n ≥ 5 for each experiment with ≥ 3 independent experiments. *P < 0.05, **P < 0.01, ***P < 0.001, Unpaired Student’s *t*-test.

### Accumulation of cytoplasmic ROS in *DEGS1*^*H132R*^ neuroblastoma

*DEGS1*^*H132R*^ is a disease-associated variant (Pant et al., 2019) which changes a conserved Histidine to Arginine in one of the putative catalytic motifs important for ceramide conversion. To explore the cellular mechanism of *DEGS1*^*H132R*^, we introduced it into the SH-SY5Y cells using the CRISPR/Cas9 knock-in system (Figure S2). To test whether *DEGS1*^*H132R*^ recapitulated the ROS accumulations seen in *ifc-KO* photoreceptors, we quantified the ROS level with staining of a superoxide probe DHE (Figures 2A-B’’). The intensity of DHE in *DEGS1*^*H132R*^ cells is significantly higher than that in the control cells (Figure 2C). Further, we transfected cyto-roGFP in the wild-type and *DEGS1*^*H132R*^ cells to examine the redox-state in the cytoplasm (Figures 2D-E’). Consistent with our findings in *ifc*-KO photoreceptors, the 405/488 nm ratio of cyto-roGFP was significantly increased in *DEGS1*^*H132R*^ mutant compared with wild-type SH-SY5Y cells (Figure 2F). Moreover, the level of mitochondrial superoxide in the *DEGS1*^*H132R*^ mutant was lower than that of wild-type control (Figures 2G-I). These data demonstrated the accumulation of cytoplasmic ROS upon DEGS1 deficiency in human neuronal cells.

**Figure 2.**
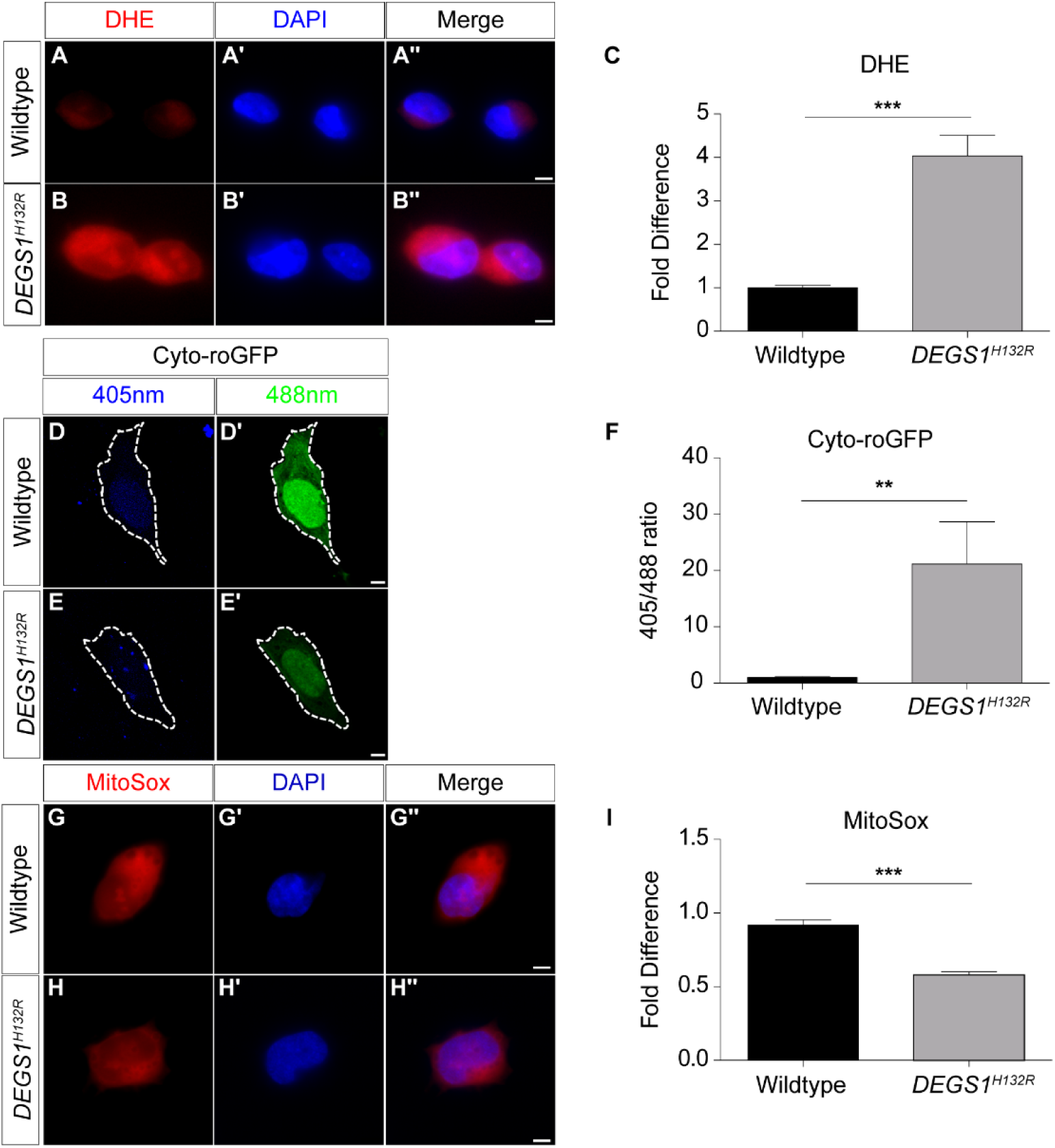
Measurement of ROS levels in CRISPR-engineered *DEGS1*^*H132R*^ SH-SY5Y neuroblastoma cells. (A-C) ROS detected in wild-type (A) and *DEGS1*^*H132R*^ (B) SH-SY5Y cells with the fluorescent probe dihydroethidium (DHE). The relative intensity of DHE was quantified in (C) with the intensity of wild-type cells set to 1. (D-F) Wild-type (D) and *DEGS1*^*H132*^ (E) SH-SY5Y cells expressing Cyto-roGFP. The region of interest for quantification is encircled by the dashed line. The 405/488 ratio of Cyto-roGFP in SH-SY5Y cells was quantified in (F). (G-I) Wild-type (G) and *DEGS1*^*H132R*^ (H) SH-SY5Y cells stained with MitoSox, a fluorescent probe specific for mitochondrial superoxide. The relative intensity of MitoSox was quantified in (I) with the intensity of wild-type cells were set to 1. Scale bars: 5 μm. Error bars represent SEM; n ≥ 3 independent experiments. *P < 0.05, **P < 0.01, ***P < 0.001, unpaired Student’s *t*-test.

### Specific NOX inhibitor, apocynin, alleviated the oxidative stress in *DEGS1*^*H132R*^ neuroblastoma

Cytoplasmic ROS are produced by cytoplasmic enzymes including NADPH oxidases (NOXs), xanthine oxidase, and uncoupled NO synthase (NOS). To identify the major ROS producer in *DEGS1*^*H132R*^ cells, we suppressed the activity of these candidate enzymes with chemical inhibition and examined the level of ROS (Figure 3G). The accumulation of ROS was ameliorated by the treatment of the NOXs inhibitor apocynin (Figures 3C-C’’) but not the xanthine oxidase inhibitor allopurinol (Figures 3E-E’’), the NOS inhibitor L-NAME (Figures 3F-F’’), or DMSO (Figures 3D-D’’; vehicle control), indicating that NOX is mainly responsible for ROS genesis in *DEGS1*^*H132R*^ cells.

**Figure 3.**
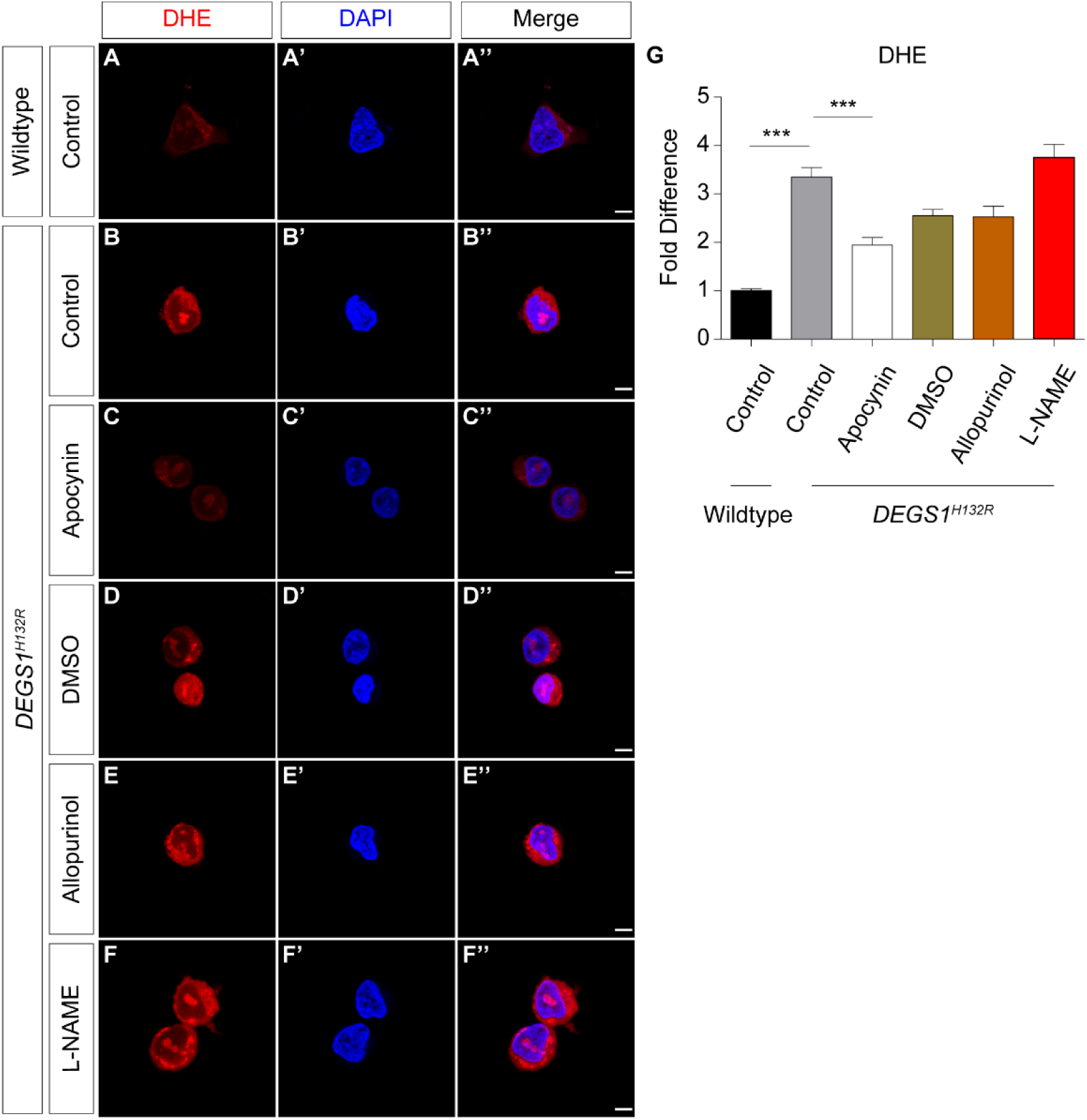
Identification of the major origins of ROS in *DEGS1*^*H132R*^ cells. (A-G) DHE staining in wild-type (A) and *DEGS1*^*H132R*^ (B) SH-SY5Y cells treated with apocynin (C) (50 μM; NADPH oxidase inhibitor) DMSO (D) (vehicle control), allopurinol (E) (100 μM; xanthine oxidase inhibitor), and L-NAME (F) (50 μM; nitric oxide synthase inhibitor) The intensity of DHE in SH-SY5Y cells was quantified in (G) with the intensity of wild-type set to 1. Scale bars: 5 μm. Error bars represent SEM; n ≥ 3 independent experiments. *P < 0.05, **P < 0.01, ***P < 0.001, one-way ANOVA with multiple comparisons.

### Activation of Rac1-NOX pathway caused the oxidative stress and subsequent neurodegeneration in *ifc*-KO photoreceptors

The small GTPase Rac1 is a cytosolic component required for NOX activation. To examine whether *DEGS1/ifc* regulates oxidative stress through Rac1 signaling, we first quantified the level of Rac1 in wild-type and *ifc*-KO eye extracts. In *ifc*-KO eyes, we saw an increase in the level of Rac1 protein compared to controls (Figures 4A-B). We then again took advantage of fly genetics and compared the activity of Rac1 in *ifc*-KO mosaic eye clones. The number of endogenous GFP-Rac1 puncta increased in the mutant clones, suggesting the activation of Rac1 (Figures 4C-D). We examined Pak1-GFP, a downstream reporter of Rac1 activation (Harden et al., 1996), and found an increase in the number of GFP puncta in *ifc*-KO photoreceptors (Figures 4E-F), consistent with the activation of the Rac1-Pak1 axis. Likewise, in *DEGS1*^*H132R*^ SH-SY5Y cells, we observed a dramatic increase in the signal of active Rac1 by immunofluorescence (Figures 4G-H). These evidence suggested that loss of *DEGS1/ifc* activates Rac1 thus may enhance its downstream NOX signaling.

**Figure 4.**
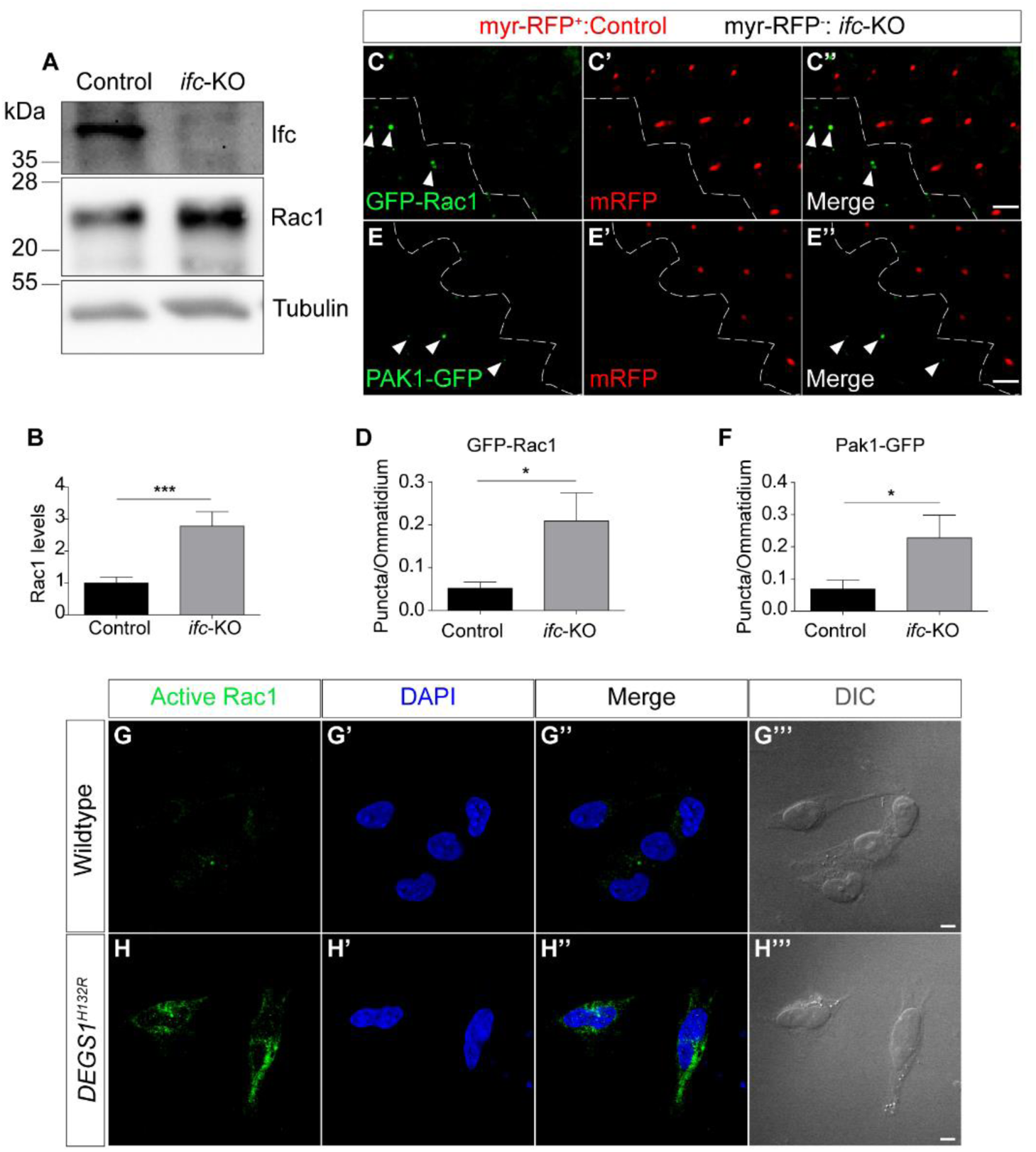
Activation of Rac1 signaling in both fly *ifc*-KO photoreceptors and SH-SY5Y *DEGS1*^*H132R*^ cells. (A-B) Western blot (A) of Rac1 comparing the eye extracts of the control and *ifc*-KO. The levels of Rac1 (normalized to tubulin) in eye extracts was quantified in (B). (C-F) Cross-section images of *ifc*-KO mosaic eye clone with endogenous expression of GFP-Rac1 (C) or PAK1-GFP (E). Control photoreceptors were marked with GMR-myr-RFP (C’, E’), separated by the dashed line from *ifc*-KO photoreceptors lacking RFP signals. Arrowheads indicate the puncta of GFP-Rac1 or PAK1-GFP. The average number of GFP-Rac1 or PAK1-GFP puncta per ommatidia was quantified in (D, F). (G-H) Immunostaining with anti-active Rac1 antibody (green) in wild-type (G) and *DEGS1*^*H132R*^ (H) SH-SY5Y cells. Scale bars: 5 μm. Error bars represent SEM; n ≥ 3 independent experiments. *P < 0.05, **P < 0.01, ***P < 0.001, unpaired Student’s *t*-test.

Next, to test whether the activation of Rac1-NOX signaling caused the degeneration in *DEGS1/ifc*-deficient conditions, we genetically suppressed Rac1 signaling in *ifc* mutant photoreceptor. The ERG amplitude of *ifc*-KO photoreceptor was restored by the genetic reduction of *rac1* (Figures 5A-B, E-F). Also, knocking-down *NOX* (Figures 5C-D, G-H) in photoreceptor or feeding flies with the NOXs inhibitor apocynin reduced the level of ROS (Figures 5K-O) and alleviated the functional decline of *ifc*-KO photoreceptors (Figures 5I-J). Thus, we concluded that the activation of the Rac1-NOX pathway increased oxidative stress and led to the subsequent degeneration of both *ifc*-KO photoreceptors and *DEGS1*^*H132R*^ neuroblastoma cells.

**Figure 5.**
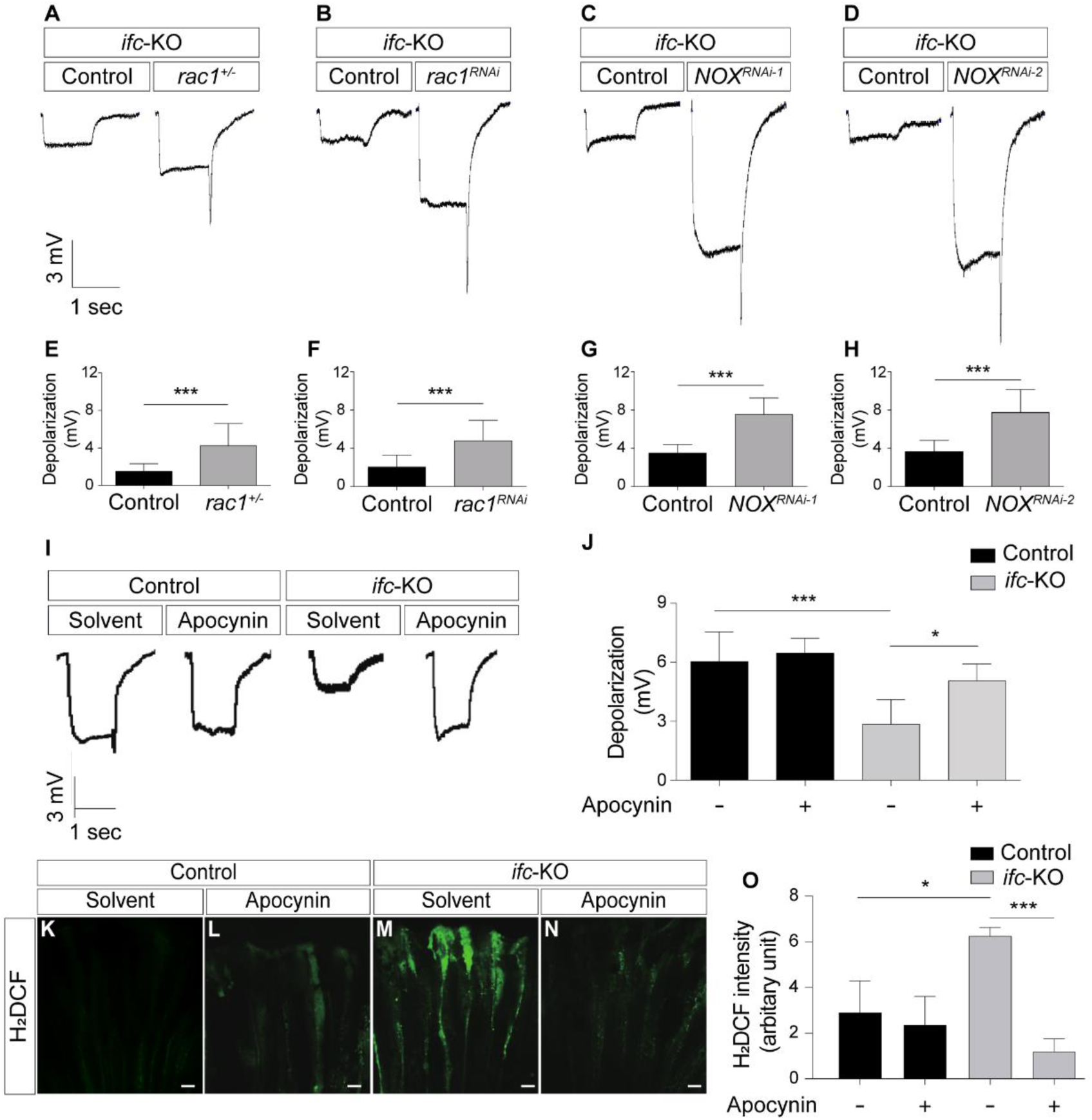
Amelioration of *ifc*-KO neuronal defects via genetic and pharmacological inhibition of the Rac1-NOX pathway. (A-H) ERG traces (A-D) upon light stimulations in *ifc*-KO eyes with genetic inhibition of the Rac1-NOX pathway after 5 days of light exposure. The UAS-*RNAi* were expressed by photoreceptor-specific *GMR*-Gal4. The average depolarization amplitude (mV) of ERG was quantified in (E-H). (I-J) ERG traces (I) of adult fly eyes with apocynin treatments and vehicle control. The average depolarization amplitude (mV) was quantified in (J). K-O H_2_DCF staining of *ifc*-KO photoreceptors longitudinal sections with apocynin treatments and controls (K-N). The ROS levels in control and *ifc*-KO photoreceptors were quantified in (O). Scale bars: 5 μm. Error bars represent SEM; n ≥ 3 independent experiments. *P < 0.05, **P < 0.01, ***P < 0.001, unpaired Student’s *t*-test.

### Loss of *ifc* increased the colocalization of active Rac1 and Rab7-positive compartments

How does *ifc* regulate the Rac1 pathway? Compartmentalization of Rac1 has been shown to play a crucial role in downstream signaling (Payapilly and Malliri, 2018). To examine whether *ifc* affects the site of Rac1 action, we examined the subcellular localization of active Rac1 in *ifc*-KO photoreceptors (Figures 6A-J’’). The colocalizations of active Rac1 puncta (arrowhead) with compartmental markers were quantified (Figures 6K-O), and we observed an increased colocalization between active Rac1 and the late endosome/lysosome marker Rab7 in *ifc*-KO photoreceptors compared with the control (Figures 6E-F’’, M). In contrast, *ifc*-KO did not affect the colocalization between active Rac1 and markers from early endosome (Rab5; Figures 6I-J’’, O), mitochondria (TOM20; Figures 6C-D’’, L), autophagosome (GABARAP; Figures 6G-H’’, N), and endoplasmic reticulum (BIP; Figures 6A-B’’, K). These data demonstrated that loss of *ifc* alters the subcellular localization of Rac1, thus suggest the critical role of ifc in mediating the site of Rac1 action for ROS production.

**Figure 6.**
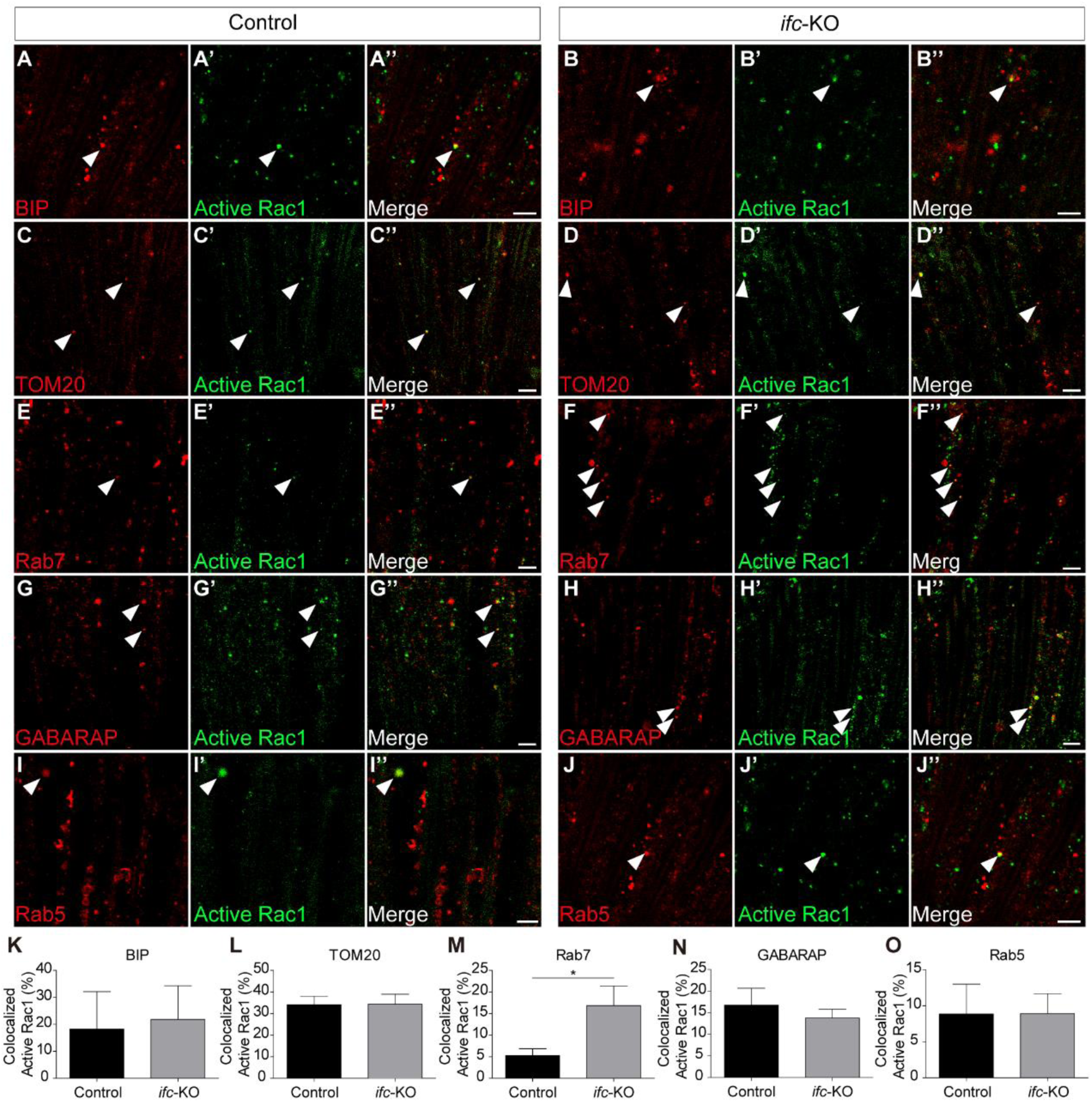
Increased subcellular colocalization of GTP-Rac1 and Rab7 in *ifc*-KO photoreceptors. (A-O) The colocalization of Active-Rac1 (green) and markers of various subcellular compartments (red) including BIP (A-B;;) (ER lumen), TOM20 (C-D’’) (mitochondrial outer membrane), Rab7 (E-F’’) (endolysosome), GABARAP (G-H’’) (autophagosome), and Rab5 (I-J’’) (early endosome) in fly photoreceptor longitudinal sections. Arrowheads indicate the colocalized puncta between GTP-Rac1 and compartmental markers. The percentage of GTP-Rac1 puncta colocalizing with compartmental markers were quantified in (K-O). Scale bars: 5 μm. Error bars represent SEM; n ≥ 3 independent experiments. *P < 0.05, **P < 0.01, ***P < 0.001, unpaired Student’s *t*-test.

### Dihydroceramide controls Rac1 association with reconstituted plasma membrane and autophagosome

Localized Rac1 signaling starts with the binding of active Rac1 to the membrane of a distinct subcellular compartment (Payapilly and Malliri, 2018). Because Ifc converts dihydroceramide to ceramide and may mediate the subcellular localization of active Rac1 (Figure 6), we hypothesized that these two sphingolipids modulate the membrane association of Rac1. To verify whether dihydroceramide and ceramide mediate Rac1 membrane binding, we performed a liposome binding assay using large unilamellar vesicles (LUVs) that mimic subcellular membranes (Figure 7). Plasma membrane raft is a well-known site of Rac1 binding; thus, we composed LUVs with the lipid compositions mimicking the membrane rafts (delPozo, 2004) (Figures 7A-C). The presence of dihydroceramide reduced the association of GTP-Rac1 (active) (Figure 7B). Next, as dihydroceramide accumulation has been shown to induce autophagy and we previously reported the activation of protective autophagy in *ifc*-KO photoreceptors (Jung et al., 2017), we tested how dihydroceramide and ceramide affect the binding of Rac1 to autophagosome-like liposomes (Rao et al., 2016) (Figures 7D-F). Strikingly, increased dihydroceramide in autophagosome-like liposome improved the membrane binding of active Rac1 (Figure 7E). In contrast, ceramide did not significantly interfere with the association of Rac1 to LUVs compared to controls, highlighting the distinct biochemical property of dihydroceramide versus ceramide in the interaction between Rac1 and membranes of different lipid compositions. These findings established a role of dihydroceramide in modulating the membrane association of Rac1.

**Figure 7.**
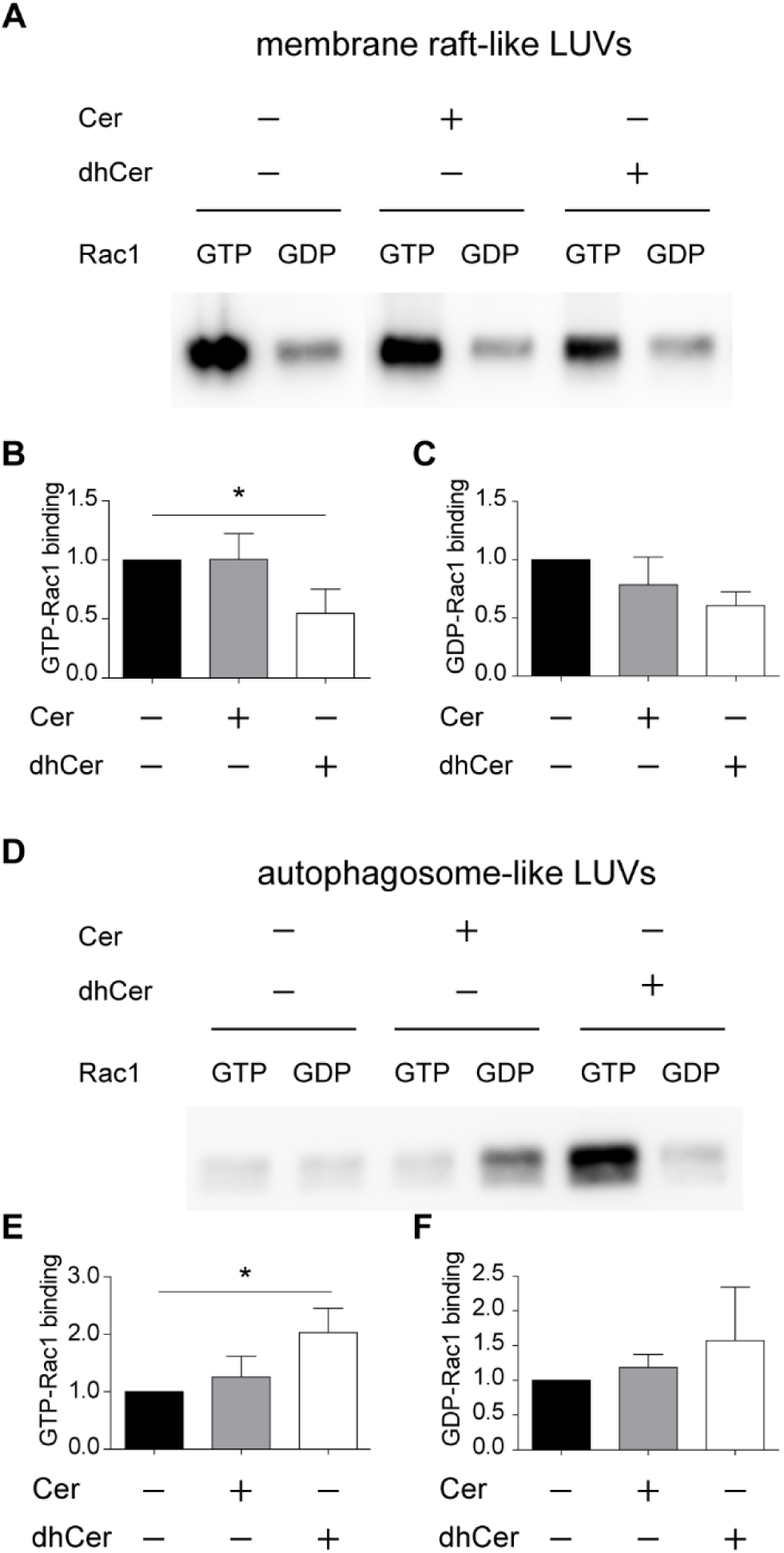
The effect of dihydroceramide on the binding of Rac1 to liposomes mimicking distinct subcellular compartments. (A-C) Western blot of Rac1 associated with membrane raft-like liposomes. The level of GTP- and GDP-Rac1 associated with membrane raft-like liposomes were quantified in (B) and (C), respectively. (D-F) Western blot of Rac1 associated with autophagosome-like liposomes. The level of GTP- and GDP-Rac1 associated with autophagosome-like liposomes were quantified in (E) and (F), respectively. Error bars represent SEM; n ≥ 3 independent experiments. *P < 0.05, **P < 0.01, ***P < 0.001, unpaired Student’s *t*-test.

## Discussion

In the present study, we investigated the subcellular origin and the molecular mechanism of ROS production in *ifc*-KO photoreceptors and *DEGS1*^*H132R*^ neuroblastoma. We found that the deficiency of *ifc* and *DEGS1* activated the Rac1-NOX complex, causing cytosolic ROS accumulation. Upon further investigation of the mechanism underlying the ROS elevation, we found that loss-of-*ifc* increased the localization of active Rac1 signaling in Rab7-positive compartments, *i.e.*, endolysosomes and autolysosomes. Finally, we demonstrated that dihydroceramide differentially mediated the binding of active Rac1 to cellular membranes. This result supported the translocation of Rac1 in the dihydroceramide-accumulating *ifc*-KO photoreceptors. In conclusion, these findings reveal a molecular mechanism connecting specific sphingolipid imbalance and oxidative signaling detrimental to neuronal maintenance.

NOX is one of the major sources of ROS, and previous studies have shown a connection between NOX-dependent oxidative stress and neural disorders (Sorce et al., 2017). However, the subcellular site of NOX action remained ambiguous. Previously, localized NOX activation has been shown to be involved in endosomal ROS signaling and endosomal NOX activation has been implicated in neurotoxic contexts (Li et al., 2011; Oakley et al., 2009). In addition, NOX-dependent ROS production from endosomes has been shown to cause neurotoxicity through NF-κB signaling in a mouse cell culture model of amyotrophic lateral sclerosis (Li et al., 2011). Besides evidence in the neuronal contexts, endosomal NOX activation may regulate VEGF signaling and promote the development of a prostate tumor cell line (Harrison et al., 2018). Knock-down of *NOX1, NOX2*, and *Rac1* abrogated c-Src and subsequent NF-κB activities from the endosomes following hypoxia/reoxygenation injury in HeLa cells (Li et al., 2008). These lines of evidence supported the significant impact downstream of NOX activation in the endosomal compartments. However, how the cytoplasmic components are recruited to the endosome remained elusive. In the present study, we demonstrated that dihydroceramide accumulation resulted in the endolysosomal activation of the Rac1-NOX complex in *ifc*-KO photoreceptors because of increased membrane binding of active Rac1. The activation of endolysosomal NOX caused the oxidative stress and degeneration in *ifc*-KO eyes. Furthermore, SH-SY5Y cells knocked-in with the disease-associated *DEGS1*^*H132R*^ variant recapitulated the Rac1-NOX activation in *ifc*-KO fly photoreceptors. Last but not the least, as apocynin treatment alleviated the oxidative damage in both systems, it indicates a specific antioxidant therapy potentially beneficial for *DEGS1*-associated neural disorders.

A growing body of literature has indicated the distinct features and pathophysiological roles of dihydroceramide in comparison to ceramide (Rodriguez-Cuenca et al., 2015; Siddique et al., 2015). In neuronal oxidative stress, no research has addressed how dihydroceramide accumulation would elevate ROS levels and the mechanisms of ceramide-induced ROS production also remained unsolved. Increased ceramide has been shown to lead to apoptosis through mitochondrial oxidative stress in neuroblastoma cells (Czubowicz and Strosznajder, 2014). Using cultured retinal neurons, exogenous ceramide was shown to decrease mitochondrial membrane potential (Prado Spalm et al., 2019). Also, NOX inhibition eased the oxidative stress resulted from TNF-α-induced ceramide production in SH-SY5Y cells (Barth et al., 2012). While these studies showed that ceramide species were cytotoxic, the findings have limited applications since these studies were performed by adding exogenous short-chain ceramides to cultured neurons, let alone the long-overlooked, less-studied dihydroceramide. To reveal the biological effects endogenous dihydroceramide, we previously knocked out *ifc*, the key gene regulating *de novo* ceramide synthesis, and demonstrated that the neurotoxicity resulted from ROS accumulation (Jung et al., 2017). We herein showed a molecular mechanism that dihydroceramide might regulate the site of Rac1-NOX activation through the association of active Rac1 to biological membranes.

The lipid composition of membranes is important for the association of Rac1 to subcellular sites, and the multi-functionality of Rac1 is accomplished by the compartmentalization (Payapilly and Malliri, 2018). For instance, phospholipids have been shown to modulate the recruitment of Rac1 to microdomains on plasma membrane (Magalhaes and Glogauer, 2010). While sphingolipids are a major class of lipid found in membrane microdomains, whether and how sphingolipids affect the site of Rac1 action was unknown. Our study examining the role of sphingolipids in Rac1-membrane interactions has highlighted dihydroceramide as a critical determinant affecting the binding of active Rac1 to membranes. As both ceramides *per se* and its enriched membrane domains have been shown to modulate translocations of a wide spectrum of proteins in distinct subcellular compartments (Fekry et al., 2018; Grassmé et al., 2001; Kajimoto et al., 2004; Zhu et al., 2019), *DEGS1/ifc*-deficiency and the consequential change in the relative levels of ceramide versus dihydroceramide may also have a broader impact on subcellular signaling in the context of neurodegeneration and beyond.

Our investigation on *ifc/DEGS1* argues that sphingolipid imbalance, especially the dihydroceramide-to-ceramide ratio, may cause neuronal defects because of altered membrane association of signaling molecules. Aside from *DEGS1*, mutations in the genes of the ceramide *de novo* synthesis pathway, including *SPTLC1, SPTLC2, CERS1*, and *CERS2*, have been associated with neural disorders in human (Bejaoui et al., 2001; Dawkins et al., 2001; Godeiro Junior et al., 2018; Karsai et al., 2019; Mosbech et al., 2014; Pant et al., 2019; Rotthier et al., 2010; Vanni et al., 2014). Nonetheless, mechanistic investigations on the neuronal defects are still in their infancy. In animal models of hereditary sensory and autonomic neuropathy Type 1 (HSAN1), ectopic expression of dominant-negative variants of the *Drosophila* and *C. elegans Spt1* gene both caused dysregulated vesicle trafficking (Cui et al., 2019; Oswald et al., 2015). In addition, a mouse model of HSAN1 with ablation of *Sptlc2* manifested significant ER-stress and prolonged activation of mTORC1 (Wu et al., 2019). Downstream of *SPTLC1/2*, deficiency in ceramide synthase 1 and 2 both caused progressive myoclonic epilepsy (Godeiro Junior et al., 2018; Mosbech et al., 2014; Vanni et al., 2014). In astrocyte culture of *CerS2* null mice, altered ceramides composition induced mitochondrial oxidative stress and impaired clathrin-mediated endocytosis (Volpert et al., 2017). As mutations of genes in the *de novo* pathway cause obvious alteration in the sphingolipidomes, the pathological role of imbalanced sphingolipids warrant detailed investigation; to this end, our findings provide an interesting and important instance of how specific sphingolipid species modulated the pathogenesis of neural disorders.

## Acknowledgements

We would like to thank Drs. Robin Hiesinger, Jennifer Jin, Ya-Wen Liu, and all members of the Chan and Huang labs for critical comments on this manuscript. We also thank Dr. Hwei-Jan Hsu, Dr. Ya-Wen Liu, Bloomington *Drosophila* Stock Center, *Drosophila* Genetic Resource Center at the Kyoto Institute of Technology, Vienna *Drosophila* Resource Center, Developmental Studies Hybridoma Bank, *Drosophila* Transgenic RNAi Project at Harvard Medical School, and the Zurich ORFeome Project for fly strains and reagents. We further thank Wellgenetics for fly injection services, the staff of the Gene Knockout/in Cell Line Modeling Core, and the Imaging Core at the First Core Labs at the National Taiwan University College of Medicine for technical assistance. This work was supported by grants from the Ministry of Science and Technology of Taiwan (MOST) (107WFA0111977) to S. -Y. H, and MOST (108-2321-B-002-060-MY2, 108-2311-B-002-011-MY3), National Taiwan University (NTU) (109L104308, 109L893603), and NTU Hospital (109C101-10-1) to C.-C. C.

## Author contributions

F.-Y. Tzou, T.-Y. Su, C.-C. Liu, S.-Y. Huang, and C.-C. Chan were involved in the conceptualization of the project. Y.-H. Yeh, Y.-L. Yu, C.-C. Liu, F.-Y. Tzou, T.-Y. Su performed the experiments and analyzed the data. F.-Y. Tzou, T.-Y. Su, S.-Y. Huang, and C.-C. Chan wrote and edited the manuscript.

## STAR Methods

### Key Resources Table

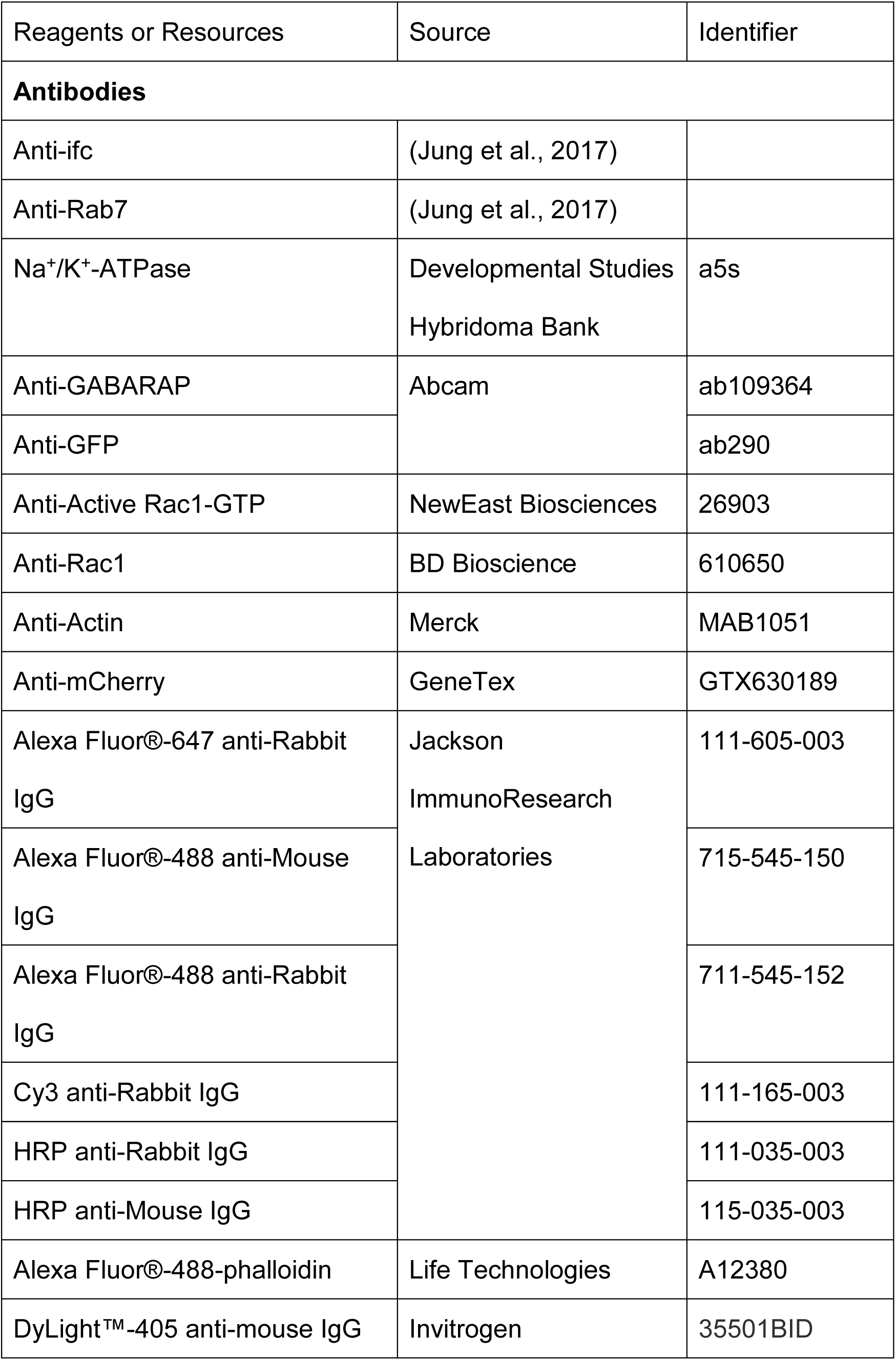
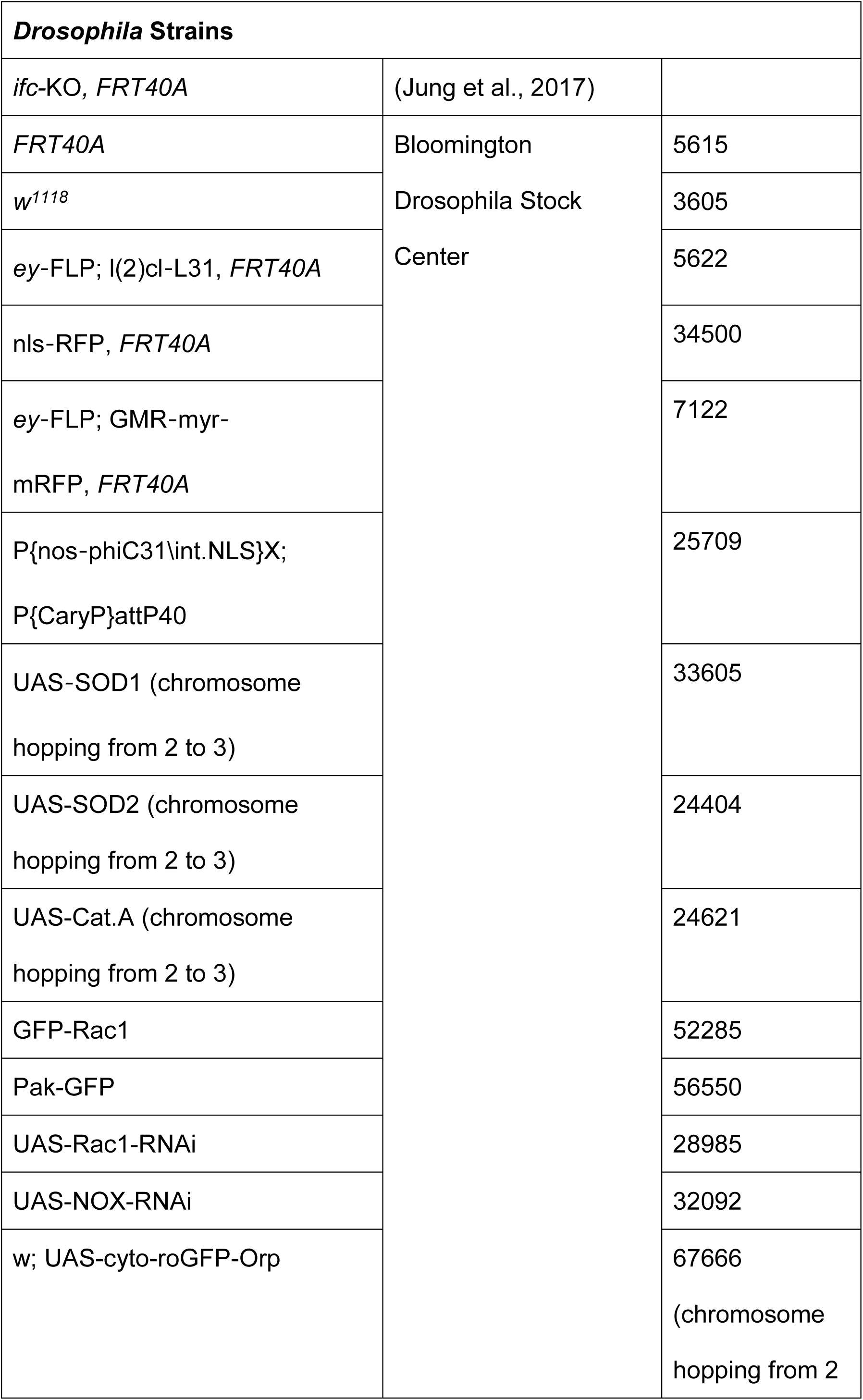
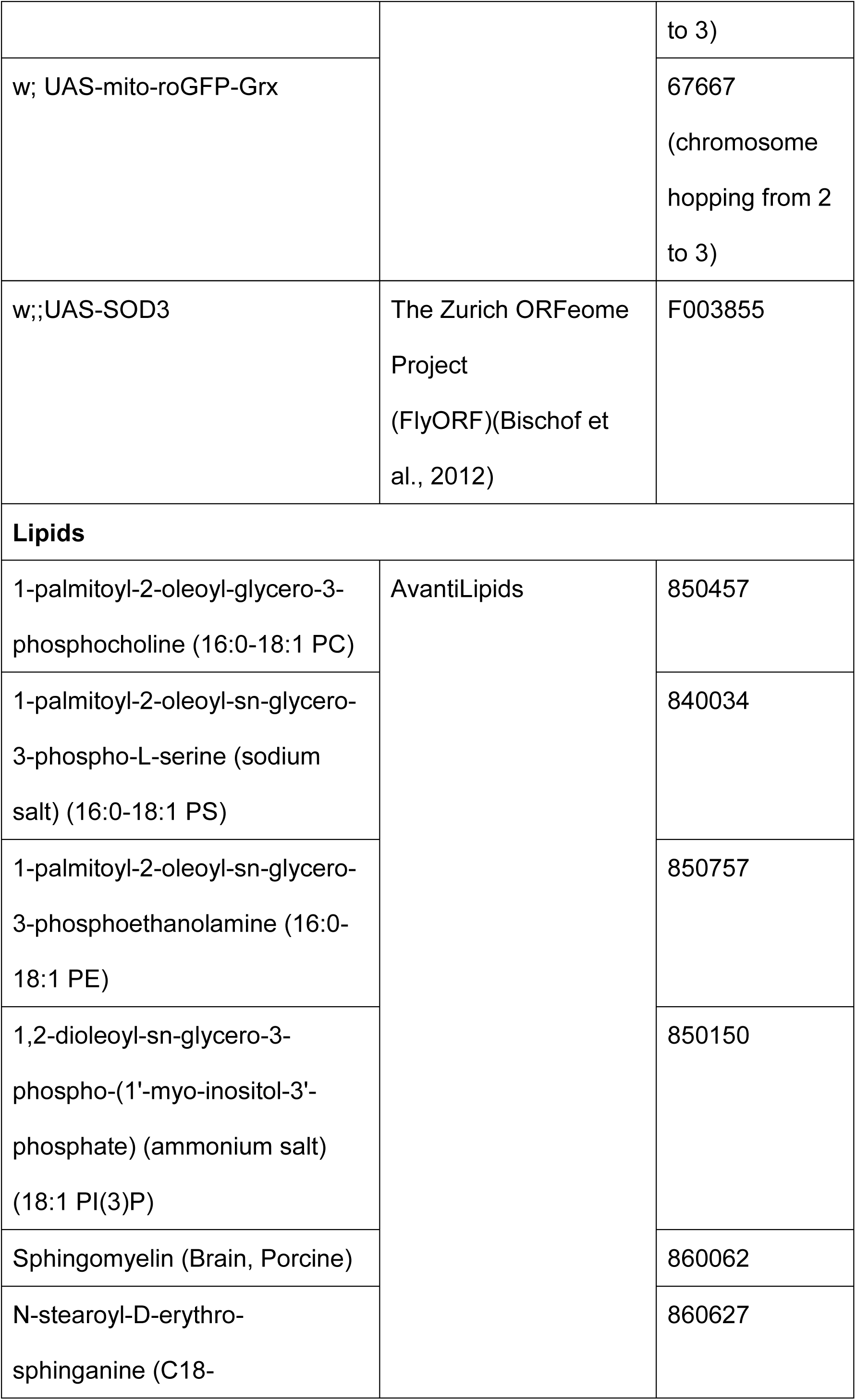
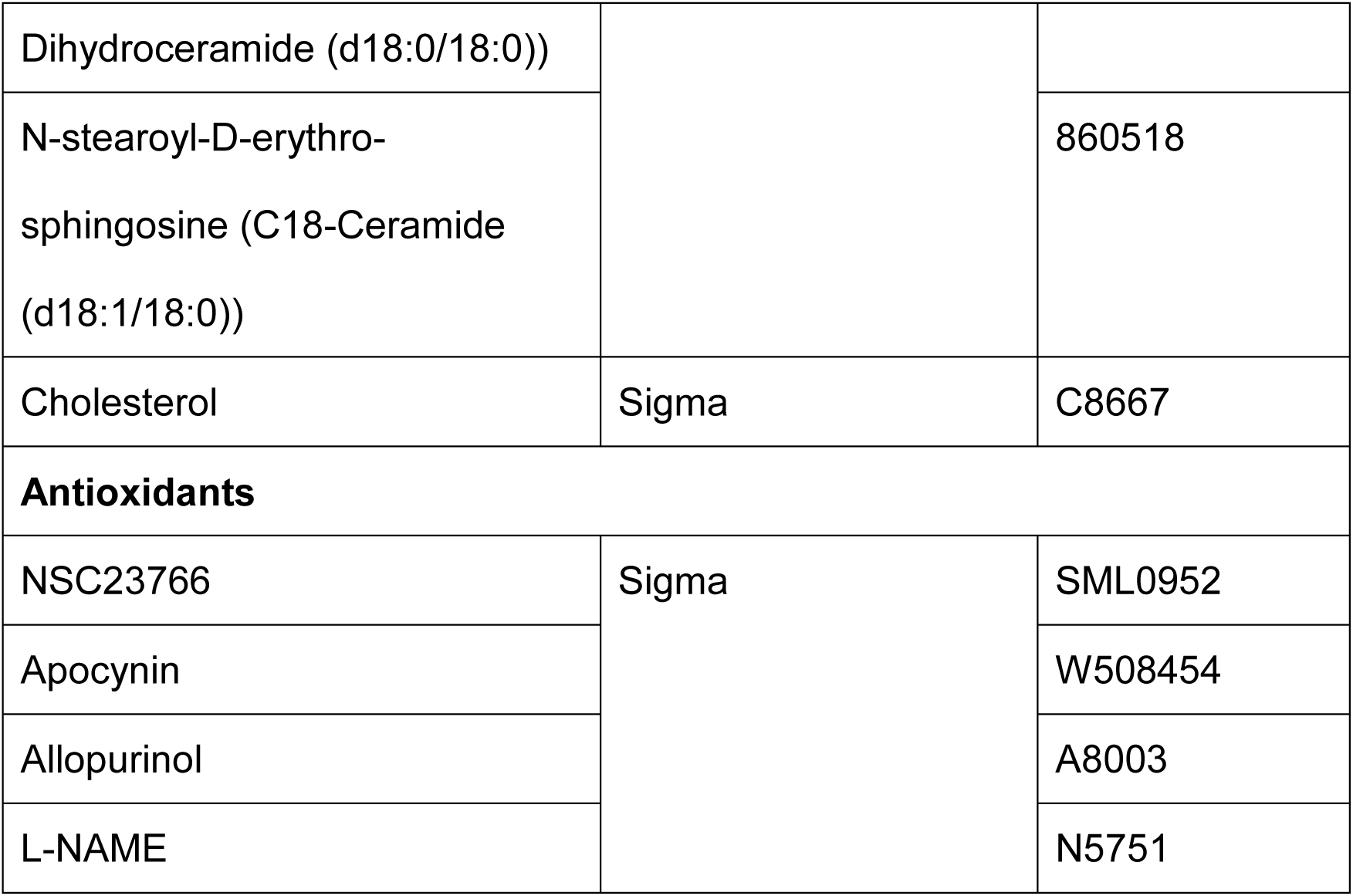

### Experimental Models and Subject Details

#### Drosophila strains and genetics

*Drosophila* stocks and crosses were maintained at 25°C on standard medium following standard fly husbandry. Information on individual fly strains can be found on FlyBase (flybase.org) unless otherwise noted (Key Resources Table).

For light stimulation, flies were exposed to constant 1000 Lux light from a LED light source. Eye-specific mosaic clones were generated using *ey-FLP* on the X chromosome while whole-eye clones of *ifc*-KO photoreceptors were generated as previously described (Jung et al., 2017).

### Method Details

#### Immunohistochemistry and microscopy

For immunofluorescence staining, adult retina tissues were dissected in Phosphate-buffered saline (PBS), fixed in 4% paraformaldehyde for 20 minutes, and washed three times for 15 minutes each in PBS with 0.4% Triton X-100 (0.4% PBST). Fixed retina tissues were incubated with primary antibodies diluted in 0.4% PBST overnight at 4°C. After primary antibody incubation, samples were washed in 0.4% PBST three times for 10 minutes, followed by secondary antibody incubation overnight at 4°C. On the next day, the samples were washed again three times in 0.4% PBST for 10 minutes before mounted in Vectashield (Vector Laboratories). ROS detection was performed with 10 μM of CM-H_2_DCFDA (chloromethyl-2′,7′-dichlorodihydrofluorescein diacetate, Thermo Fisher, C6827) following the manufacturer’s protocol. Samples were analyzed on either a Leica TCS SP5 confocal microscope with LAS AF software or a Zeiss LSM 880 confocal microscope with Zen software. Imaging data were processed and quantified using the Fiji package of ImageJ (National Institutes of Health) (Schindelin et al., 2012).

##### Electroretinogram (ERG)

ERGs were performed as previously described (Jung et al., 2017). In brief, flies were immobilized on glass slides, and the ERGs were recorded upon one second light pulses with the recording and reference electrodes filled with 2 M NaCl. All experiments were carried out in triplicates with minimally 10 recordings per replicate for each genotype and experimental condition.

##### roGFP reporter assays

roGFP fly strains obtained from the Bloomington Drosophila stock center (Key Resources Table) were recombined into *ifc*-KO background, and mosaic clones were generated to compare the roGFP activity between the control and *ifc*-KO clones. After light exposure for 5 days, the flies were dissected following the aforementioned immunohistochemistry protocol. The oxidation of the cyto-roGFP alters the excitation spectrum from 488 nm to 405 nm. The 405/488 ratio was used as an index for redox state.

##### Western blotting

Protein extracts were separated by SDS-PAGE and transferred to polyvinylidene difluoride membranes (PVDF, Millipore, IPVH00010, pore size: 0.45 μm) as per manufacturer’s instructions (Bio-Rad). PVDF membranes were incubated with 5% non-fat milk in TBST [10 mM Tris (pH 8.0), 150 mM NaCl, 0.1% Tween 20] for 1 hour at room temperature, washed three times with TBST for 10 minutes, and incubated with primary antibodies at 4°C overnight. Membranes were washed three times with TBST for 10 minutes before incubation with secondary antibodies in TBST for 1 hour at room temperature. Blots were then washed with TBST three times, developed with ECL reagents (Thermo, 34580; Millipore, WBKLS0500; or GE health, RPN2235), and captured by the BioSpectrum™600 Imaging System (UVP Ltd).

##### Rac1-Liposome Binding Assay

For liposome preparations, lipid mixtures were dried and rehydrated in Buffer 1 [10 mM HEPES (pH 7.5), 140 mM NaCl, and 1.5 mM MgCl_2_]. The lipid compositions for liposomes mimicking membrane-raft and autophagosome are listed below in percentage, and the total concentration of lipid mixture was 1 mM.

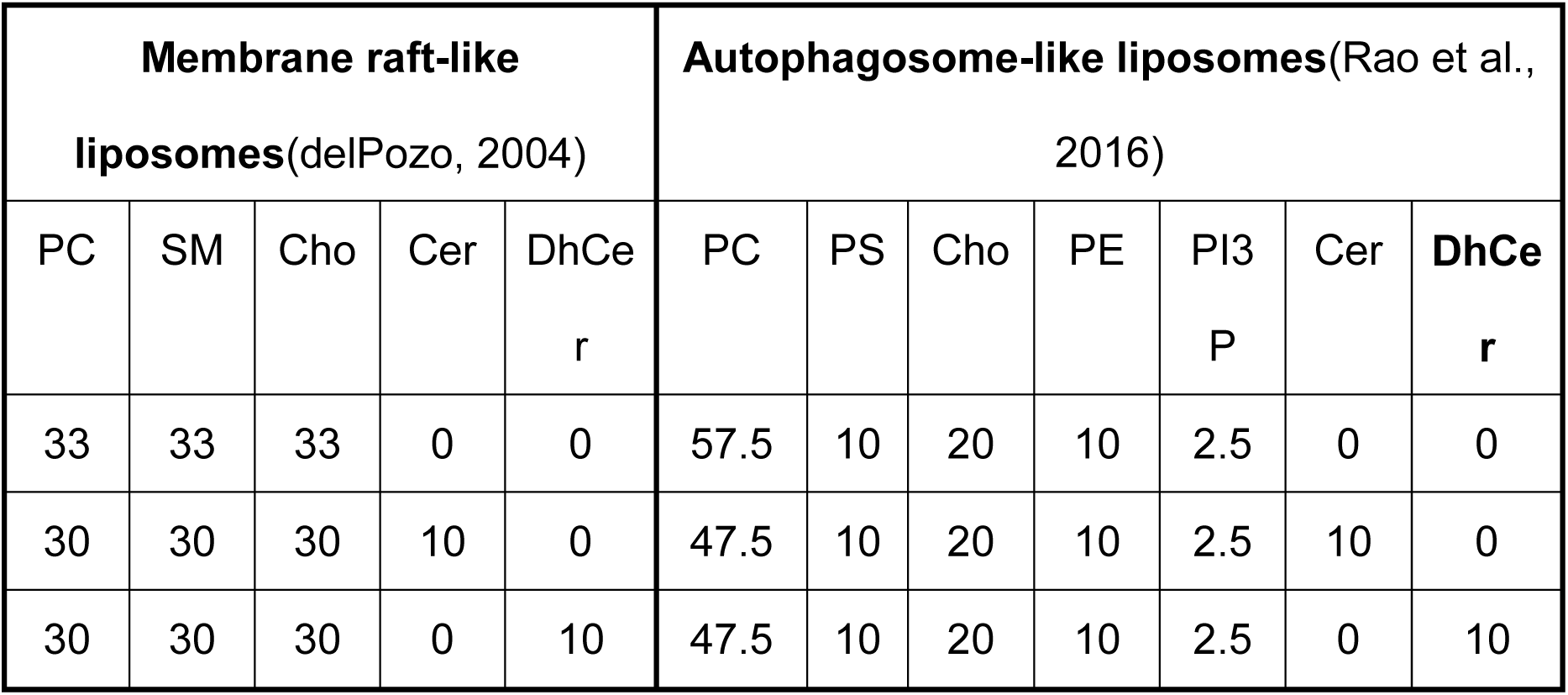

Next, the liposome mixtures were subjected to a series of freeze-thaw cycles before being extruded through 0.1 μm pore size polycarbonate membranes (Whatman, Cat. 800309) using Avanti Mini-Extruder. C18 ceramide and dihydroceramide were used since C18 ceramides are the predominant species in the neuronal context (Laviad et al., 2008).

Recombinant Rac1 (Abcam, ab51014) was incubated with 1x GTPγS (Sigma, 20-176) or GDP (Sigma, 20177) in Buffer 2 [40 mM HEPES (pH7.5), 4 mM EDTA, 2 mM DTT] for 10 minutes at 30°C. The resultant protein mixtures were prenylated by rabbit reticulocyte lysate (Promega, L415A) and 2 μM farnesyl pyrophosphate (Sigma, F6892) in Buffer 3 (50 mM Tris, pH 7.5, 5 mM MgCl_2_, 50 μM ZnCl_2_ 2 mM DTT) for 30 min at 37°C.

To determine membrane binding, prenylated Rac1 (1 μg) was added to 100 μl of 1 mM liposomes and incubated for 30 minutes at 37°C. Prenylated Rac1 protein was separated into soluble and membrane-bound fractions by 40,000 x g, 15-minute centrifugation at 4°C. Liposome pellets were washed 3 times with Buffer 1 before heating at 95°C for 10 min with 50 μl of 1x Laemmli Sample Buffer for Western blotting.

##### DEGS1 CRISPR knock-in SH-SY5Y cell

Human *DEGS1*^*H132R*^ knock-in point mutation was generated using the CRISPR/Cas9 system. The sgRNA design for CRISPR/Cas9 target sites was computationally identified at Zhang Lab’s website (http://crispr.mit.edu/) and cloned by using the all-in-one plasmid construction Kit with dual selections of puromycin resistance and fluorescence (pZG22C02, ZGene Biotech Inc.). Trypsinized SH-SY5Y cells were washed with PBS once, and 1 × 10^5^ cells mix with 2 µg plasmids plus 40 pmole single strand donor were used per electroporation (1100v/50ms/1pulses) using Neon™Transfection System 10 µl Kit (MPK1025, Thermo Fisher Scientific). The cells were plated into 24-well plates containing antibiotic-free complete medium and incubated at 37°C overnight in a 5% CO2 incubator before the selection with 0.5µg/ml puromycin for 2 days. A subset of the cells was harvested for the analysis of editing efficiencies, and the rest were diluted to 0.5 cells per well and plated into 96-well plates for single-cell cultures for at least one month. Half of the cells in each well were lysed for genomic PCR and the PCR products were treated with ExoSAP PCR clean-up reagent (75001, Thermo Fisher Scientific) and then send for sequencing. When the sequence results showed a pattern consistent with the target mutation, the PCR products were then cloned into a TA-cloning vector for blue-white selection. A total of 10 white colonies were selected and subjected to PCR amplification with M13 forward and reverse primers. The PCR products were cleaned and send for sequence confirmation. The genomic PCR for detecting *DEGS1*^*H132R*^ knock-in SH-SY5Y was performed using the following primer set:

Forward primer: 5’-TTTGGGGCCTATGCGTTTGG-3’

Reverse primer: 5’-GTGCAAACCCAGGCCAAGTA-3’

##### Cell Culture, DHE staining, and measurement of neurite length

The wild-type and *DEGS1*^*H132R*^ knock-in SH-SY5Y human neuroblastoma cells were cultured in DMEM/F12 medium (Gibco, 12500062) supplemented with 10% fetal bovine serum (Gibco, A3160502) and maintained at 37°C in a 5% CO_2_ humidified incubator. The cells were passaged when reaching confluency, usually every 7 days.

For Dihydroethidium (DHE; Invitrogen, D11347) staining, cells were seeded on cover glasses (Glaswarenfabrik Karl Hecht GmbH, 41001118) in 12-well plates at a density of 1 × 10^5^ cells per well. The wild-type or *DEGS1*^*H132R*^ knock-in SH-SY5Y cells were treated with DMSO, 100 μM Trolox, 100 μM mito-TEMPO, 100 μM apocynin, 100 μM Allopurinol, and 100 μM L-NAME for 24 h before incubating with 5 μM DHE in PBS (Gibco, 10010-023) for 30 min at 37°C. Stained cells were washed with PBS and fixed with 4% paraformaldehyde (Electron Microscopy Sciences, 15710) in PBS for 10 min at room temperature. Fixed cells were washed with PBS, stained with 300 nM DAPI (Invitrogen, D1306) in PBS for 5 min at room temperature, and washed again with PBS. All samples were mounted in Vectashield (Vector Laboratories) and analyzed on a Zeiss Axio Imager M1 or LSM880 microscope.

For the measurement of neurite length, the wild-type and *DEGS1*^*H132R*^ knock-in SH-SY5Y cells were seeded on 6-well plates at a density of 1 × 10^5^ cells/well and incubated overnight. Neural induction was initiated the following day by adding fresh medium containing 10uM retinoic acid (RA). The cells were imaged every 24 hours for 3 days, and culture media with 10 μM RA was changed every other day. Multiple images were taken under light microscopy and the captured images were labeled with a scale bar and quantified using Fiji (Schindelin et al., 2012).

### Quantification and Statistical Analysis

Quantitative data were analyzed using two-tailed unpaired Student’s *t*-test or one-way ANOVA with Tukey’s multiple comparison test and graphs were generated using GraphPad Prism 5. All data in bar graphs were expressed as mean ± SEM. P-values of less than 0.05 were considered statistically significant.

## Supplemental Information titles and legends

**Supplemental Figure 1.**
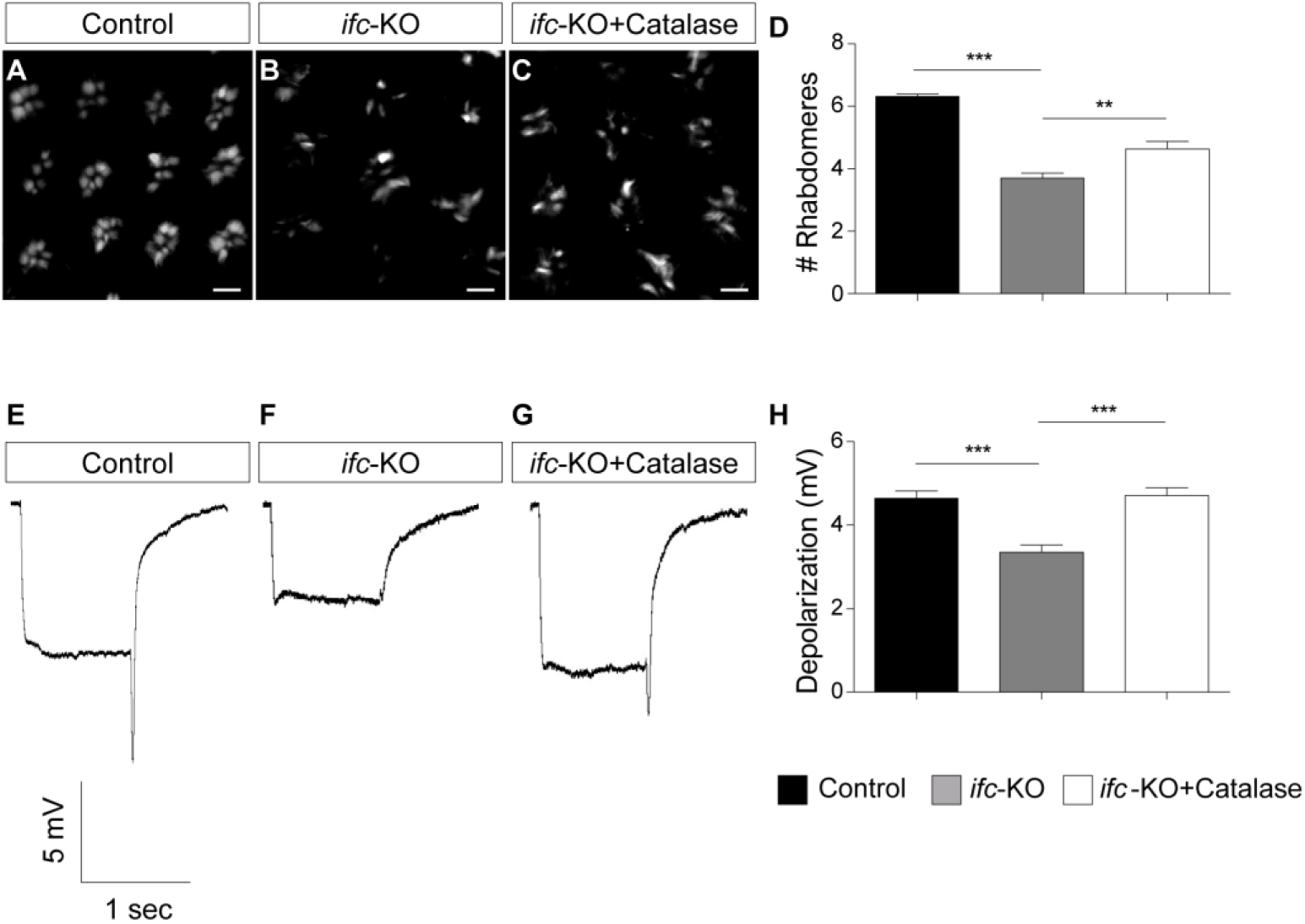
Catalase overexpression ameliorates the neuronal defects in *ifc*-KO. (A-D) Photoreceptors cross sections stained with Phalloidin (white) after 5 days of light exposure. The average number of rhabdomeres per ommatidia was quantified in (D). (E-H) ERG traces of flies after 5 days of light exposure. Double-ended arrows indicate the depolarization amplitude. The average depolarization amplitudes (mV) of adult fly eyes was quantified in (H). Scale bars: 5 μm. Error bars represent SEM; n ≥ 3 independent experiments. *P < 0.05, **P < 0.01, ***P < 0.001, unpaired Student’s *t-*test.

**Supplemental Figure 2.**
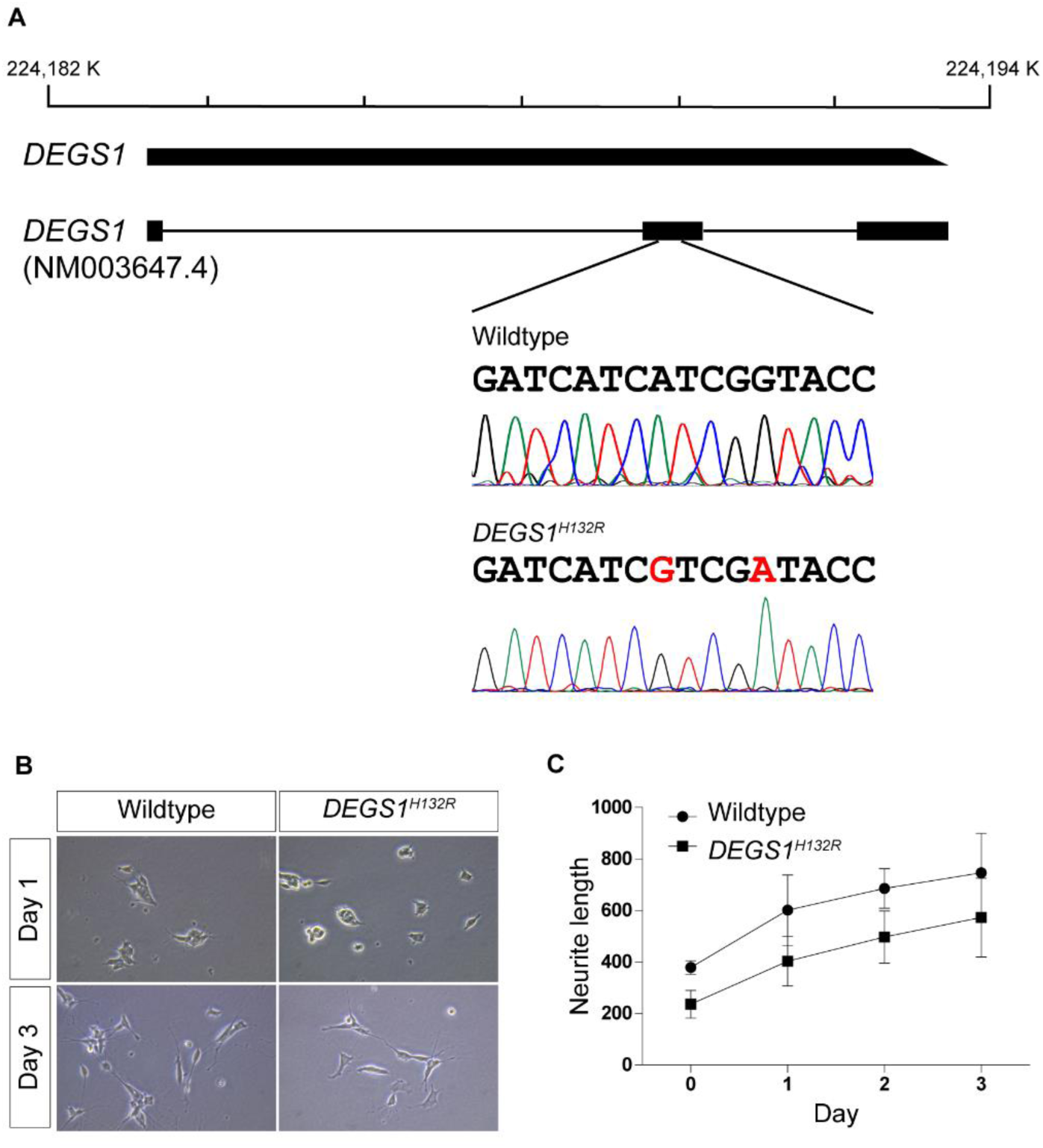
CRISPR knock-in of *DEGS1*^*H132R*^ in SH-SY5Y neuroblastoma cells. (A) A schematic diagram of *DEGS1*^*H132R*^ CRISPR knock-in. (B-C) Phase contrast images of wild-type and *DEGS1*^*H132R*^ SH-SY5Y cells with retinoic acid (RA, 10 μM) induction for 1 day or 3 days. The average neurite length of RA-treated wild-type and *DEGS1*^*H132R*^ SH-SY5Y was quantified in (C). Error bars represent SEM; n ≥ 3 independent experiments. *P < 0.05, **P < 0.01, ***P < 0.001, unpaired Student’s *t-*test.

